# Oviposition status and sugar availability influence measures of transmission potential in malaria-infected mosquitoes

**DOI:** 10.1101/2024.01.25.577126

**Authors:** Justine C. Shiau, Nathan Garcia-Diaz, Dennis E. Kyle, Ashutosh K. Pathak

**Author notes:** Corresponding author. Justine C Shiau. Nathan Garcia-Diaz, Dennis E Kyle.

## Abstract

**Background:** Like other oviparous organisms, the gonotrophic cycle of mosquitoes is not complete until they have selected a suitable habitat to oviposit. In addition to the evolutionary constraints associated with selective oviposition behavior, the physiological demands relative to an organism’s oviposition status also influences their nutrient requirement from the environment. Yet, studies that measure transmission potential (vectorial capacity or competence) of mosquito-borne parasites rarely consider if the rates of parasite replication and development could be influenced by these constraints resulting from whether mosquitoes have completed their gonotrophic cycle.

**Methods:** *Anopheles stephensi* mosquitoes were infected with *Plasmodium berghei* the rodent analog of human malaria and maintained on 1% or 10% dextrose and either provided oviposition sites (‘oviposited’ herein) to complete their gonotrophic cycle or forced to retain eggs (‘non-oviposited’). Transmission potential in the four groups was measured up to 27 days post-infection as 1) the rates of vector survival, 2) rates of sporozoite migration to the salivary glands (‘extrinsic incubation period’ or EIP), and 3), sporozoite densities.

**Results:** In the two groups of oviposited mosquitoes, rates of sporozoite migration and densities in the salivary glands were clearly higher in mosquitoes fed 10% dextrose. In non-oviposited mosquitoes however, rates of sporozoite migration and densities were independent of sugar concentrations, although both measures were slightly lower than oviposited mosquitoes fed 10% dextrose. Rates of vector survival were higher in non-oviposited mosquitoes.

**Conclusions:** Taken together, these results suggest vectorial capacity for malaria parasites may be dependent on nutrient availability and oviposition/gonotrophic status and as such, argue for more careful consideration of this interaction when estimating transmission potential; costs to parasite fitness and vector survival were buffered against changes in nutritional availability from the environment in non-oviposited mosquitoes, but not oviposited mosquitoes. In general, however, these patterns suggest parasite fitness may be dependent on complex interactions between physiological (nutrition) and evolutionary (egg-retention) trade-offs in the vector, with potential implications for disease transmission and management. For instance, while reducing availability of oviposition sites and environmental sources of nutrition are key components of integrated vector management strategies, their abundance and distribution is under strong selection pressure from the patterns associated with climate change.

**Graphical abstract:** 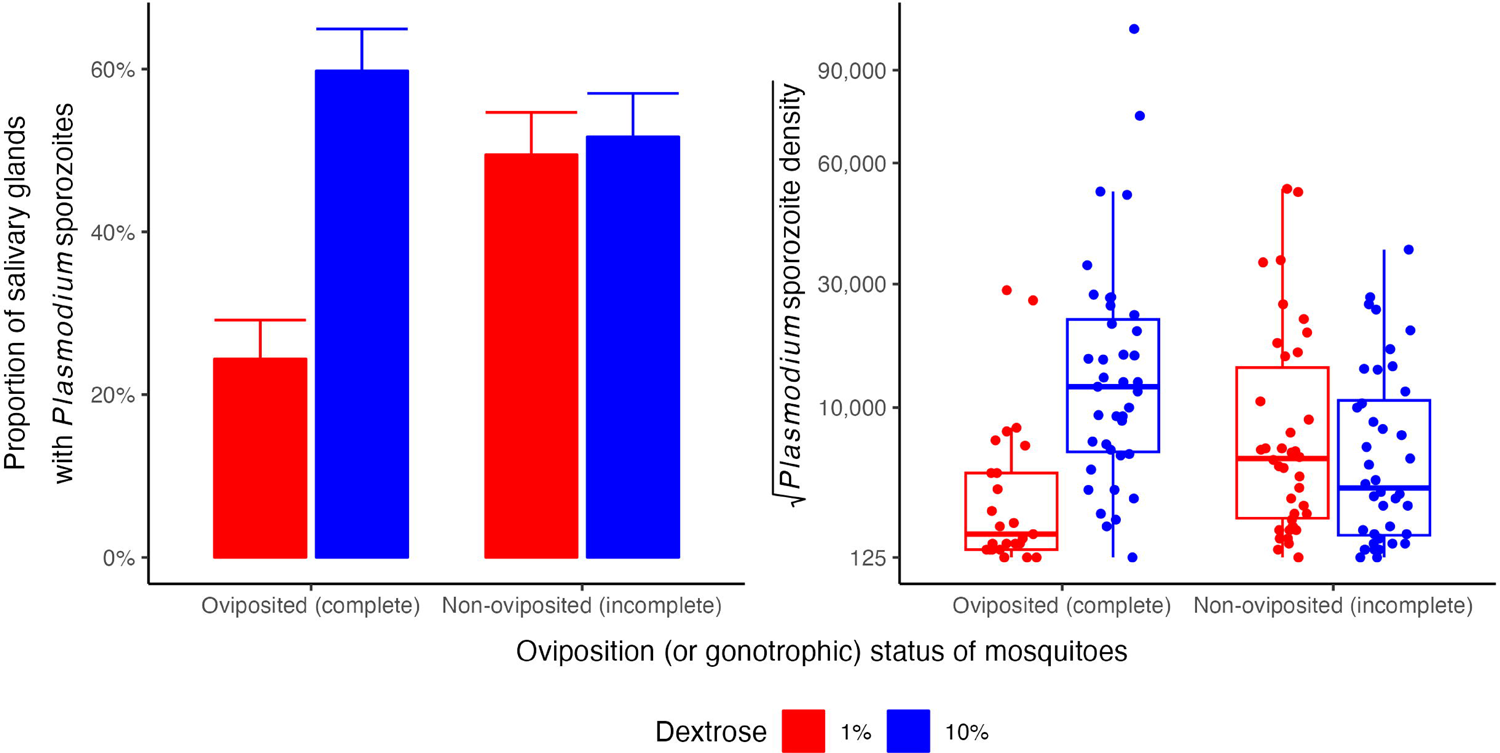

## INTRODUCTION

Like other oviparous organisms, mosquitoes retain their eggs until they find a suitable habitat to oviposit and complete its gonotrophic cycle. Amongst oviparous organisms however, mosquitoes occupy an important niche: most species rely exclusively on hematophagy (blood feeding) of vertebrates for producing eggs, which enables them to vector several parasites with devastating consequences for the health and fitness of humans and wildlife. Following ingestion of an infected blood meal, egg and parasite development may be initiated at the same time in the mosquito. However, by the time a mosquito has completed egg development, parasite development may or may not be complete, although transmission to the next host is dependent on the mosquito locating an oviposition site to complete its ‘gonotrophic cycle’ [1]. While several studies suggest widespread changes in mosquito behavior and physiology associated with the gonotrophic cycle [2, 3, 4, 5, 6, 7, 8], few have determined if and how changes in oviposition status influences parasite development rates and transmission potential [9]; in general, measures of parasite fitness and transmission potential are performed in mosquitoes prevented from completing their gonotrophic cycle (e.g., see [10, 11, 12, 13] and references therein). A greater understanding of how parasite fitness is shaped by oviposition status of mosquitoes is important for at least two reasons. First, integrated vector management strategies either directly or indirectly alter the availability of oviposition sites, by managing water sources for instance, to identifying specific physical, chemical, and biological cues underlying selective oviposition behavior for targeting with chemical or biological means [1, 14, 15, 16, 17, 18, 19]. Second, the availability of oviposition sites will continue to change as a cumulative response to climatic factors such as global warming and changes in rainfall patterns to the increasing urbanization and deforestation [20, 21, 22, 23, 24]. By altering the number and distribution of oviposition habitats, both scenarios could select for variation in oviposition behavior and thus modify parasite/disease transmission in the process.

Like other oviparous organisms, adult, female mosquitoes also show selective oviposition behavior [1, 16]. Offspring survival and reproduction are benefited by ovipositing mothers choosing sites that fulfil several criteria, including for instance, habitat size, quality, risk of predation and presence of conspecifics. The ubiquity of selective oviposition behavior attests to its benefits to future reproduction, with selection for this trait imposed at the cost of reduced fecundity for instance [25, 26, 27, 28, 29, 30]. While several studies have shown how variation in this behavior could modify the oviparous organism’s own fitness [26, 27, 28, 30, 31, 32, 33, 34, 35], few have assessed whether the associated trade-offs could have downstream effects on the fitness of other organisms such as parasites, whose life-history strategy is driven by its interaction with the oviparous host.

Organismal fitness is also dependent on how it allocates available resources from the environment to meet the costs of various life-history traits [25, 26, 27, 29, 36]. For vector-transmitted parasites, although its growth and reproduction are dependent on how they allocate resources from the mosquito host, the host’s own fitness is also dependent on how they allocate resources from environmental sources of nutrition; in other words, nutritional availability from the environment can shape life-history strategies of the vector and parasite [37, 38]. Amongst environmental sources of nutrition, the effect of sugars on life-history traits of adult, female mosquitoes are particularly well characterized, with their propensity for sugar feeding also exploited for controlling vector density (e.g. toxic sugar baits) [39]; while the costs of parasite infection for the mosquito are more apparent at lower concentrations of sugar, in general, sugar requirements are greater in gravid/non-oviposited mosquitoes foraging for oviposition sites, carrying and/or maintaining eggs [25, 38, 40, 41, 42]. Differences in sugar concentrations and/or availability to the mosquito have also been shown to affect some parasite life-history traits, especially for *Anopheles* mosquitoes infected with the unicellular, eukaryotic microparasites of the genus *Plasmodium* [38]; these parasites are the etiological agent of malaria, a disease that continues to extract significant costs in morbidity and mortality in humans [43]. In malaria-infected mosquitoes, *Plasmodium* oocyst densities in the midguts and mosquito survival were dependent on dextrose (‘D-glucose’) concentrations [44]. Taken together, these observations suggest that differences in nutritional requirements between oviposited and non-oviposited mosquitoes could influence parasite fitness.

In principle, a single *Plasmodium* parasite ingested with a blood meal can invade the midgut wall of an adult, female *Anopheles* and differentiate into an oocyst. Within the oocyst, in a process termed ‘schizogony’, several rounds of asexual replication culminate in the generation of sporozoites. Once mature, the oocysts rupture and sporozoites are released into the mosquito hemocoel, from where they migrate to the salivary glands, ready for injection during the next blood meal. In general, transmission to the next host (i.e., future reproduction) is dependent on three traits: 1) the vector surviving past the time required for sporozoites to invade the salivary glands, 2) rates of sporozoite migration to the salivary gland (‘extrinsic incubation period’, EIP herein) [45, 46], and 3), the number of sporozoites in the salivary glands [47]. Although a single oocyst can produce thousands of sporozoites, all three fitness traits (vector survival, rates of migration, and the overall densities) are dependent on trade-offs with other traits in the parasite and vector’s life history strategy [45, 48, 49, 50, 51]. The current study tests the hypothesis that parasite fitness would be dependent not only on whether the mosquito has oviposited or not (oviposition status), but also the amount of nutrition available. Female *An. stephensi* mosquitoes were exposed to an infectious blood meal containing *P. berghei* [49, 50]. Vector survival, rates of sporozoite migration and densities were measured in mosquitoes that were (or not) allowed to oviposit, and whether this was affected by differences in nutritional availability (1% or 10% dextrose) [40].

## MATERIALS AND METHODS

All reagents and consumables were purchased from Thermo-Fisher Scientific, unless stated otherwise.

### Study design

For a visual overview of the study design, refer to Additional file 1: Fig. S1. Based on procedures described in detail previously [49, 50], 1500 adult, female *An. stephensi* were infected with *P. berghei* infected mice (0 days post-infection (dpi herein)). At day 1 post-infection, ∼350 mosquitoes were transferred to each of 4 cages for the respective treatments. Two cages were provided a low nutrition diet (1% dextrose), while the remaining two were offered the high nutrition diet (10% dextrose). At 6 dpi, 1 cage from each nutrient treatment were provided oviposition sites until 9 dpi [50]. The overall study lasted 27 days and employed a fully crossed design wherein the same starting population of *P. berghei*-infected mosquitoes were separated across four treatment groups: 1) oviposited mosquitoes with low nutrient availability (1% dextrose), 2) oviposited mosquitoes with high nutrient availability (10% dextrose), 3) non-oviposited mosquitoes with low nutrient availability and finally, 4) non-oviposited mosquitoes with high nutrient availability (Additional file 1: Fig. S1A).

Vector mortality, rates of *P. berghei* sporozoite migration (referred to herein as EIP) and sporozoite densities in the salivary glands were quantified essentially as described previously [49, 50]. While vector mortality and EIP are well known higher order components of the vectorial capacity model [45, 46], sporozoite densities in the salivary glands are increasingly recognized as a critical fitness-conferring trait in the next vertebrate host [47]. Mosquito mortality was monitored daily in all 4 cages from day 1 post-infection [50]. To determine EIP and overall sporozoite prevalence [50], salivary glands from 15-20 mosquitoes from each group were checked for presence or absence of sporozoites at three-day intervals starting 9 dpi (i.e., days 9, 12, 15, 18, 21, 24 and 27 post-infection). To enumerate sporozoite densities [49], salivary glands from 10-15 mosquitoes from each group were sampled from days 15 to 27 post-infection, also at three-day intervals (i.e., 15, 18, 21, 24 and 27 dpi); note that due to differences in assay procedures (see below), EIP [50] and sporozoite densities [49] were obtained from different mosquitoes. Finally, to determine whether mosquitoes had oviposited or not, ovaries of each sampled mosquito was assessed for presence (non-oviposited) or absence (oviposited) of eggs (Additional file 1: Fig. S1B).

### Mosquito husbandry

*An. stephensi* mosquitoes (Walter Reed Army Institute of Research ca 2015, “Indian” strain) were maintained in a level 2 Arthropod containment laboratory at the University of Georgia, as described previously [52]. Colonies were maintained at 27°C ± 0.5°C, 80% ± 5% relative humidity, and under a 12-hour day/night photoperiod. Female mosquitos were provided with human whole blood (Interstate Blood-Bank, Memphis, TN) with glass membrane feeders (Chemglass Life Sciences, NJ) to support egg production. Hatched larvae were dispensed at a density of 300 L1s/1000 ml of water and maintained on pelleted Hikari cichlid gold diet (HikariUSA, Hayward, CA). Adult colonies were maintained on sugar water composed of 5% dextrose (w/v) and 0.05% para-aminobenzoic acid (PABA) (w/v).

### *P. berghei* infections of mice

All animal procedures described herein were performed as per AUP number A2020 01-013-Y3-A10, approved by UGA IACUC. Four-to six-week-old male, Hsd:ICR(CD-1) mice (Envigo, Indianapolis, IN) were infected with *P. berghei* GFP-LUC_CON_, a transgenic strain expressing GFP and Luciferase reporter genes in an ANKA background (referred to herein as *P. berghei*), as described previously [49]. Briefly, 7 mice were infected with a dose of 5×10^6^ parasites/ml in a 500µl inoculum. At 4 dpi, mice were anesthetized using 1.8% tri-bromo ethanol (Avertin, Sigma-Aldrich, MO) just prior to mosquito feeds.

### *P. berghei* infections of mosquitoes

At three-to-seven-days post-emergence, host-seeking, female mosquitoes were sorted into a cage (32 cm^3^, Bug Dorm, Taiwan) and transferred to a 20°C environmental chamber (Percival Scientific, Perry, IA). As described previously [49], anesthetized *P. berghei*-infected mice were placed atop the cage. Mosquitoes were allowed to feed on the mice for 15 minutes (day 0 post-infection). At 24 hours post-infection (or day 1), mosquitoes were distributed into 4 separate cages and maintained for the duration of the experiment as described above (‘Study design’). Mosquitoes were maintained on either 1% or 10% dextrose (w/v) supplemented with 0.05% para-Aminobenzoic acid (w/v).

### Data collection

While daily rates of mortality were recorded for all 4 treatments, starting 9 dpi and up to 27 dpi, the salivary glands of mosquitoes from each group were checked for the presence or absence of sporozoites as described previously [50]. Briefly, mosquitoes were aspirated directly into 70% ethanol (v/v) before salivary glands were dissected from the mosquito and transferred to a 3μl drop of PBS on a clean glass slide. Salivary glands were ruptured to release sporozoites by placing a cover glass directly on the drop of PBS; presence or absence of sporozoites was assessed on an upright DIC microscope (Leica DM2500) at 100x or 400x magnification. Salivary glands were sampled from 15 to 17 mosquitoes in each group at days 9, 12, 15, 18, 21, 24 and 27 post-infection.

Sporozoite densities in the salivary glands were enumerated as described previously [49]. Briefly, at days 15, 12, 18, 21, 24 and 27 post-infection, ten mosquitoes from each group were terminally sampled by aspirating directly into 70% ethanol. Salivary glands from each mosquito were resuspended in 50μl of PBS supplemented with 0.1% bovine serum in a 0.65ml tube. Sporozoites were released by homogenizing the salivary glands with 10 strokes of a microtube pestle. Sporozoites were quantified in a 10μl aliquot of the homogenate as described previously [49]; while this approach can detect ≥125 sporozoites, the relatively laborious and time-consuming nature of this assay meant quantification of sporozoite densities was not initiated until 15 dpi when appreciable (and thus more reliable) quantities of sporozoites would be available for enumeration.

### Data analyses

All data analyses were performed with generalized linear models (GLM) in JMP Pro software (ver 16). In general, prevalence data were modeled with a binomial distribution, and count data were modeled with a type 2 negative binomial distribution. For sporozoite prevalence and sporozoite densities, time was standardized and treated as a continuous fixed effect. Oviposition status and nutrient levels (1% vs 10% dextrose) were treated as categorical fixed effects. The effect of time was also modeled as a quadratic effect (‘humped’) to account for saturation and/or decline in the dependent variable trends. Gravid status (presence or absence of eggs) was analyzed with the same fixed effects described above using a binomial distribution. Mosquito survival was analyzed using the Cox Proportional Hazards model with the interaction between oviposition status and nutrient levels as fixed effects.

The analyses also considered the possibility that not all mosquitoes in the respective groups may respond to the two oviposition treatments, and as such, could potentially confound analyses and comparisons between groups. For instance, while some mosquitoes may still retain eggs despite being offered oviposition sites, in the groups not offered an oviposition site, some mosquitoes may nonetheless oviposit; assessing the contribution of these confounders is also relevant to the vector survival data where it was not possible to quantify oviposition status post-hoc [50]. To address this, mean sporozoite prevalence or mean sporozoites densities from all four groups at each time point were quantified from all mosquitoes irrespective of whether they responded to the treatment, Pearson’s product-moment correlations were performed with the mean sporozoite prevalence and densities from the same groups, but after excluding non-responder mosquitoes from the respective groups, i.e., by excluding non-oviposited (or gravid) mosquitoes from the groups offered oviposition site, or oviposited mosquitoes (not gravid) from the groups not offered oviposition sites.

## RESULTS

### Data and treatment summary

For the duration of the study, a total of 616 *An. stephensi* mosquitoes were sampled from all four groups, with 416 sampled for EIP and 200 for sporozoite densities. To determine the effectiveness of the oviposition treatments, gravid status from each mosquito (presence or absence of eggs) was assessed at the time of dissection. In the groups that *were offered* oviposition sites (‘oviposited’ herein), 89.1% of mosquitoes successfully laid eggs (or 271 of 304 mosquitoes dissected), with 90.5% (133 of 147 mosquitoes) and 87.9% (138 of 157 mosquitoes) oviposited mosquitoes recovered from low (1% dextrose) and high (10% dextrose) nutrition groups, respectively, over the duration of the experiment (Additional file 2: Fig. S2). While similar number of oviposited mosquitoes were recovered at each time point (Chi-squared test (*χ^2^*) = 0.02, *df* = 1, *P* = 0.88), recovery was also not affected by nutritional availability over time (*χ^2^* = 0.22, *df* = 1, *P* = 0.64) or nutritional availability in general (*χ^2^* = 0.49, *df* = 1, *P* = 0.48) (Additional file 3: Table S1).

In the groups that were *not offered* an oviposition site (‘non-oviposited’ herein) (Additional file 2: Fig. S2), 78.5% of the mosquitoes still retained eggs in their ovaries (or 245 of 312 mosquito ovaries assessed), with the 84.8% of mosquitoes retaining eggs in the groups offered low nutrition (134 of 158 mosquitoes) higher than the 72.1% in the high nutrition group (111 of 154 mosquitoes) (*χ^2^* = 6.86, *df* = 1, *P* = 0.008) (Additional file 3: Table S1). However, this difference was due to the apparent decline at day 24 post-infection (right pane, Additional file 2: Fig. S2A): excluding this sampling point from the analysis suggested that while nutrient availability did not affect recovery of non-oviposited mosquitoes (*χ^2^* = 0, *df* = 1, *P* = 0.9686), similar numbers of non-oviposited mosquitoes were recovered at the chosen sampling times (*χ^2^* = 0.22, *df* = 1, *P* = 0.6322). As the reason for this discrepancy was unknown, all statistical analyses were performed with this sampling point included in the dataset.

Despite the presence of these mosquitoes in the respective groups (i.e., oviposited individuals in groups not offered an oviposition cup, or non-oviposited individuals in groups offered oviposition cups), correlation analysis suggested that in general, these individuals should not confound downstream data analyses (Additional file 4: Fig. S3): compared to the estimations of mean sporozoite prevalence and densities from all four groups over time irrespective of oviposition status, excluding these individuals did not significantly alter the estimation of prevalence (Pearson’s correlation coefficient, *r_(26)_* = 0.99, *P*<0.001) (Additional file 4: Fig. S3A), or sporozoite densities (Pearson’s *r_(18)_*= 0.96, *P*<0.001) (Additional file 4: Fig. S3B). Thus, all analyses below were performed with datasets that consider all individuals irrespective of whether they responded to the oviposition sites as expected; furthermore, this approach also ensures compatibility with the vector survival data where it was not possible to discriminate between mosquitoes that may or may not have responded as expected to the oviposition sites.

### Time to infectiousness (EIP) and overall sporozoite prevalence in salivary glands is dependent on nutritional availability in mosquitoes that have oviposited, but not those that retain eggs

Of the 616 mosquitoes, salivary glands from a total of 416 mosquitoes from all four groups were assessed for the presence or absence of sporozoites at 9, 12, 15, 18, 21, 24 and 27 dpi (Fig. 1). Differences in nutritional availability to oviposited and non-oviposited mosquitoes affected the rates at which the mosquitoes became infectious with sporozoites in the salivary glands (EIP) (*χ^2^* = 5.45, *df* = 1, *P* = 0.02) (row 8, Table 1) (Fig. 1A), as well as the overall proportion of sporozoite-positive salivary glands (*χ^2^* = 5.45, *df* = 1, *P* = 0.02) (row 5, Table 1) (Fig. 1B). This effect is particularly apparent in oviposited individuals that were maintained at 1% dextrose (left pane, Fig. 1A, and Fig. 1B). Indeed, pairwise comparisons of the marginal means predicted by the model suggested a consistently lower prevalence in this group compared to the other three groups after 15 dpi, with differences in estimated probabilities predicted to range from −0.55 to −0.23 (rows 1-21, Additional file 5: Table S2). While nutritional availability did not appear to result in clear differences between non-oviposited mosquitoes, for the groups maintained at 10% dextrose, mosquitoes were infectious earlier in the oviposited group, with higher prevalence at 15 dpi (Estimated difference in probability (*Est.* herein) = 0.34, *standard error (SE)* = 0.1, *P* = 0.004) and to a lesser yet significant extent at 18 dpi (*Est.* = 0.27, *SE* = 0.1, *P* = 0.048) (rows 38 and 39, Additional file 5: Table S2).

**Fig. 1.**
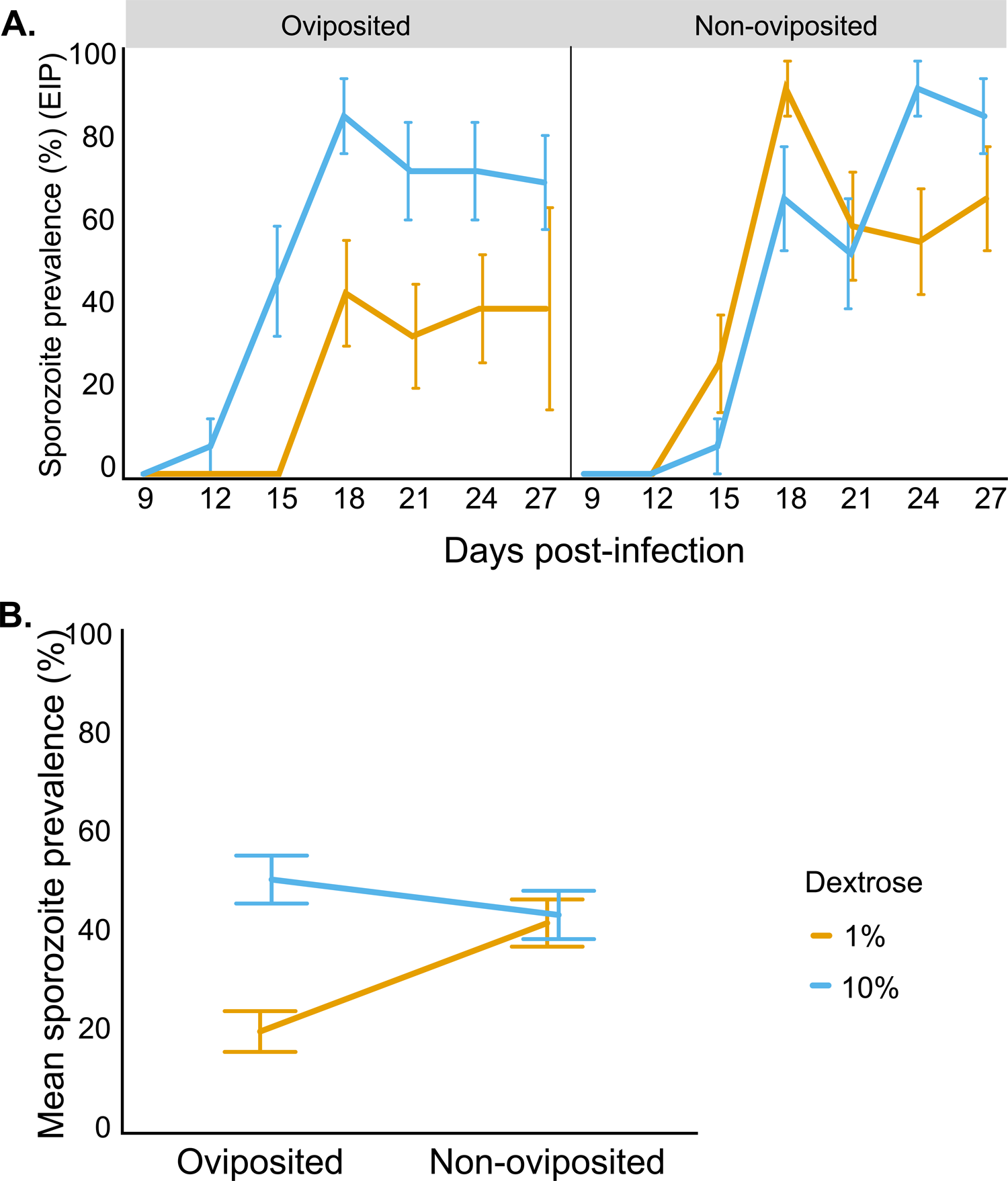
**A)** Time from infection till mosquitoes became infectious (‘extrinsic incubation period’ or EIP) based on appearance of sporozoites in the salivary glands of oviposited (left pane) and non-oviposited (right pane) mosquitoes, maintained on a diet of low (1% dextrose, yellow) or high nutrition (10% dextrose). **B)** Mean sporozoite prevalence aggregated over the time points in A). Refer to Table 1 for statistical modeling of the data set, and Additional file 5: Table S2 for post hoc pairwise comparisons of the four groups at each day post-infection.

**Table 1.** Statistical modeling of sporozoite prevalence (‘*y* = sporozoite prevalence’) (Figure 1) and sporozoite densities (‘*y* = sporozoite densities’) (Figure 2). Binomial and negative binomial generalized linear models respectively, were used to model sporozoite prevalence and densities.

| Row | Terms (*x*) | *y* = Sporozoite prevalence |  |  | *y* = Sporozoite densities |  |  |
| --- | --- | --- | --- | --- | --- | --- | --- |
| | | $\chi^2$ | *df* | *P* | $\chi^2$ | *df* | *P* |
| 1 | Oviposition status <sup>a</sup> | 3.14 | 1 | 0.076 | 0.69 | 1 | 0.404 |
| 2 | Nutrient levels <sup>b</sup> | 12.37 | 1 | <0.001* | 6.95 | 1 | 0.008* |
| 3 | Dpi <sup>c</sup> (linear effect) | 33.94 | 1 | <0.001* | 4.29 | 1 | 0.038* |
| 4 | Dpi (humped/‘quadratic’ effect) | 31.95 | 1 | <0.001* | 1.78 | 1 | 0.183 |
| 5 | Oviposition status * nutrient levels | 10.67 | 1 | 0.001* | 12.94 | 1 | <0.001* |
| 6 | Oviposition status * dpi | 2.86 | 1 | 0.091 | 1.22 | 1 | 0.269 |
| 7 | Nutrient levels * dpi | 1.87 | 1 | 0.171 | 2.50 | 1 | 0.114 |
| 8 | Oviposition status * nutrient levels<br>* dpi | 5.45 | 1 | 0.02* | 5.36 | 1 | 0.021* |
|  | Number of salivary glands | 416 |  |  | 142 (200 sampled) |  |  |

<sup>a</sup> Oviposited or Non-oviposited; <sup>b</sup> 1 or 10% dextrose; <sup>c</sup> salivary glands sampled at 3-day intervals from 9-27 dpi for sporozoite rates, and 15-27 dpi for densities.
Abbreviations: $\chi^2$ = Chi-squared test (type 2); *df* = degrees of freedom; dpi = days post-infection; *P* = p-value.
\* Statistical significance set at $P \leq 0.05$ .

The dynamics of sporozoite migration and overall prevalence were generally higher in oviposited mosquitoes maintained at 10% dextrose compared to the other groups, albeit not as clearly as the oviposited groups maintained on 1% dextrose (Additional file 5: Table S2). In other words, the low prevalence in the 1% dextrose group was compensated by the high prevalence in the 10% dextrose group such that the mean prevalence in oviposited groups (mean = 36.8%) were comparable to the non-oviposited groups (mean = 43.4%), which could explain why oviposition status alone did not clearly influence prevalence (*χ^2^* = 3.14, *df* = 1, *P* = 0.076) (Table 1). In contrast, the clear effect of nutrition on prevalence (*χ^2^* =12.37, *df* = 1, *P* = 0.008) (Table 1) was likely driven by the strong reduction in prevalence in the oviposited group maintained on 1% dextrose, which, despite the higher prevalence in the non-oviposited mosquitoes also maintained on 1% dextrose, resulted in mean prevalence of 32.2% that was lower than the mean of the groups maintained at 10% dextrose (47.4%). Finally, the analyses also suggested that when averaged over all four groups, the initial increase in proportion of sporozoite-positive salivary glands recovered (*χ^2^* = 33.94, *df* = 1, *P*<0.001) was eventually followed by saturation and/or decline (*χ^2^* = 31.95, *df* = 1, *P*<0.001) (Table 1) after 18 dpi (Fig. 1A). Taken together, these results suggest complex consequences of interactions between nutrient availability and oviposition status of mosquitoes on sporozoite migration rates.

### Sporozoite densities in the salivary glands are dependent on nutritional availability in mosquitoes that have oviposited, but not those that retain eggs

Of the 613 mosquitoes sampled in total, 200 salivary glands were assessed for sporozoite densities at 15, 18, 21, 24 and 27 dpi (Fig. 2); 142 individuals showed evidence of carrying ≥125 sporozoites (see “Methods”). In general, the trends were similar to sporozoite prevalence, with differences in nutritional availability to oviposited and non-oviposited mosquitoes affected the dynamics of sporozoites densities in infected salivary glands (*χ^2^* = 5.36, *df* = 1, *P* = 0.021) (row 8, Table 1) (Fig. 2A), as well as overall sporozoite densities (*χ^2^* = 12.94, *df* = 1, *P*<0.001) (row 5, Table 1) (Fig. 2B). Pairwise comparisons of the marginal means suggested lower sporozoite densities in oviposited groups maintained at 1% dextrose compared to the other three groups (rows 1-15, Additional file 6: Table S3). In general, while sporozoite densities increased more rapidly in oviposited mosquitoes maintained on 10% dextrose, overall densities in the salivary glands were also higher compared to the other groups (Additional file 6: Table S3).

**Fig. 2.**
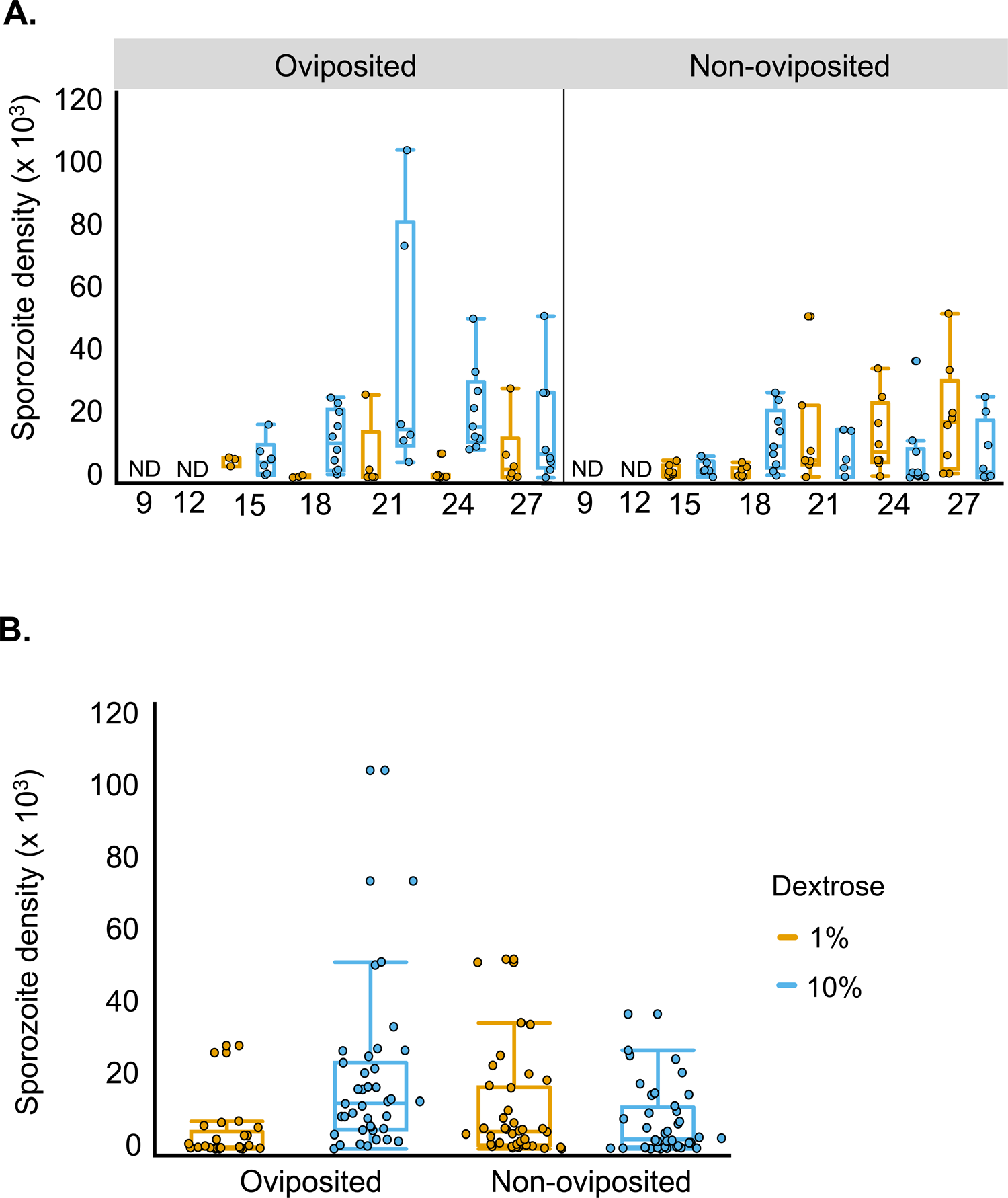
**A)** Dynamics of sporozoite densities in the salivary glands of oviposited (left pane) and non-oviposited (right pane) mosquitoes (infected mosquitoes only), maintained on a diet of low (1% dextrose, yellow) or high nutrition (10% dextrose). **B)** Mean sporozoite densities aggregated over the time points in A. Refer to Table 1 for statistical modeling of the data set, and Additional file 6: Table S3 for post hoc pairwise comparisons of the four groups at each day post-infection. The minimal limit of detection with this technique is 125 sporozoites. ND = not done (for rationale, refer to Study design under “Methods”).

### Survival is reduced in mosquitoes that have oviposited, but also due to low nutrition

Survival was dependent on oviposition status (*χ^2^* = 15.37, *df* = 1, *P*<0.001) and to a lesser extent on nutritional concentrations (*χ^2^* = 4.94, *df* = 1, *P* = 0.026) (Figure 3) (Additional file 7: Table S4), with higher risk of mortality to mosquitoes that had already oviposited (risk ratio (*rr*) = 1.42, *SE* = 0.13, *P*<0.001) and maintained at 1% dextrose (*rr* = 1.23, *SE* = 0.11, *P* = 0.025). However, mosquito survival rates were not affected by the combination of nutrient concentrations and oviposition status (*χ^2^* = 2.36, *df* = 1, *P* = 0.124) (Additional file 7: Table S4).

**Fig. 3.**
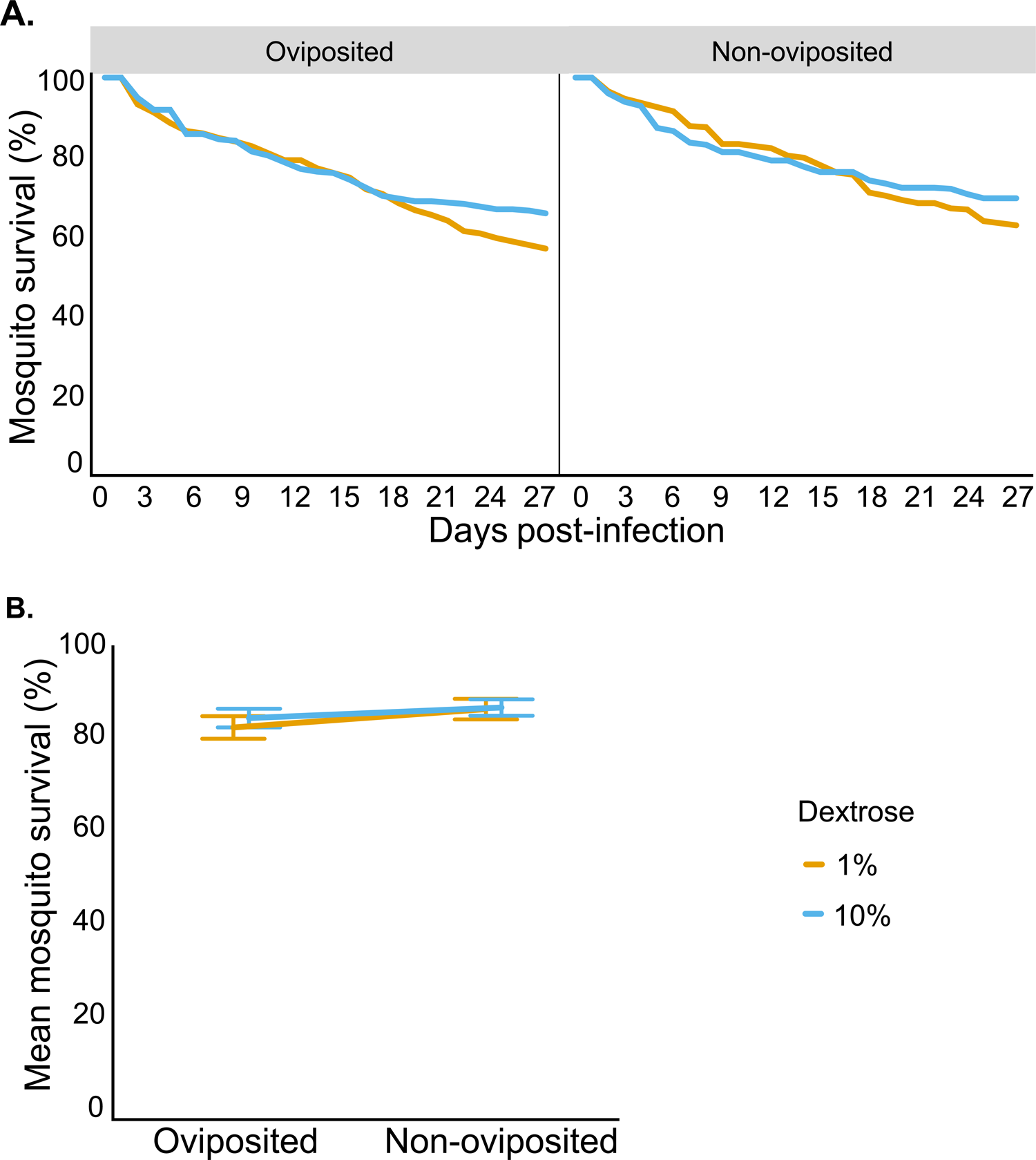
**A)** Survival rates of oviposited (left pane) and non-oviposited (right pane) mosquitoes (infected mosquitoes only), maintained on a diet of low (1% dextrose, yellow) or high nutrition (10% dextrose). **B)** Mean survival rates aggregated over the time points in A).

## DISCUSSION

Nutrient availability and oviposition status of mosquitoes resulted in complex consequences for *Plasmodium* fitness traits. Differences in nutrient availability from the environment influenced the rates at which oviposited and non-oviposited became infectious (EIP), sporozoite densities in salivary glands, and vector survival. In general, however, nutritional availability had less of an impact on the three traits in mosquitoes that were prevented from ovipositing. Taken together, these results suggest that costs to parasite fitness may be ‘buffered’ by tradeoffs experienced by mosquitoes carrying eggs, even in a resource-poor environment. However, for assays measuring transmission potential, where mosquitoes are generally not allowed to oviposit, both oviposition status and nutritional availability should receive critical consideration. For instance, at the very least, depending on oviposition status and/or nutritional availability, transmission potential may either be underestimated or overestimated.

The positive effect of environmental nutrition on parasite fitness (i.e., probability of transmission to the next host) was clearly evident in the mosquito groups that were allowed to oviposit: while higher sugar concentrations enhanced rates of sporozoite migration (Fig. 1A, left pane) and densities (Fig. 2A, left pane) in the salivary glands, higher vector survival (Fig. 3A, left pane), especially after 15 days, indicate a larger proportion of transmission-capable mosquitoes. In general, higher nutrient levels enhance sporozoite migration and densities by increasing the rates of replication of sporozoites within the oocysts lining the basal lamina of the mosquito’s midgut [53]. While low glucose concentrations adversely affected the survival of adult mosquitoes in general [40, 41], *An. gambiae* infected with *P. falciparum* showed increased attraction to and uptake of sugars, presumably in response to the higher energetic demands [38, 54, 55]. Despite the potential for compensatory feeding behavior, the concentrations used in the current study (1% glucose) were likely too low to prevent the apparent decline in mosquito survival in the oviposited mosquitoes after 15-18 dpi [40]. Although mounting immune defenses is energetically costly in insects, it could explain reduced sporozoite rates and densities in this group, with associated pathology resulting in reduced vector survival [28]. Clearly, determining whether these costs to parasite fitness are due to a general lack of resources or how the host allocates resources or energetic budgets is critical [25, 32].

Nutrition did not have such a strong influence on parasite fitness in the mosquito groups that were not allowed to oviposit (right panes in Fig. 1A, Fig. 2A and Fig. 3A). This effect was especially apparent when comparing the non-oviposited individuals maintained on a low nutrient diet with their oviposited counterparts. Indeed, despite the low nutrition, overall rates of sporozoite migration (Fig. 1B) and densities (Fig. 2B) were comparable to the high nutrient groups, with no clear costs to vector survival (Fig. 3B). While this indicates alternative sources of nutrition were able to support sporozoite replication and vector survival, reduced immune defenses due to reproduction-immunity tradeoffs in non-oviposited mosquitoes may also result in the within-host conditions being more conducive to parasite and vector survival [28]. In the absence and/or reduced availability of environmental sources of nutrition, energetic demands in adult stages of holometabolous insects can be met by the nutrients stored in the fat bodies, but also by resorption of nutrients from reproductive tissues (also referred to as ‘follicular resorption’, ‘oosorption’ or ‘follicular atresia’), albeit at the expense of reduced fecundity [32, 56, 57, 58, 59]. Since the current study was undertaken with mosquitoes originating from a common pool (Additional file 1: Fig. S1), the potential contribution of the fat bodies should be similar between the four treatment groups; in other words, all else being equal, the difference in rates of parasite replication and vector survival between the non-oviposited and oviposited groups maintained under low nutrient conditions should, in principle, reflect the extent of nutrient resorption from reproductive tissues.

In mosquitoes, nutrient resorption follows apoptotic death of ovarian follicles (referred to as follicular resorption herein) [57, 58, 60]. Follicular resorption is a trait that is shared across several orders of insects and allows the organism to adjust to physiological demands [56]. While the trait incurs costs in reduced fecundity for the current generation, evolutionary theory suggests these costs are traded off with enhanced life span, which in turn allows the insect more time to locate oviposition sites that can benefit the fitness of future generations [25, 27, 30, 31, 32, 34, 35, 58]. In the current study, survival was indeed higher in the non-oviposited group and particularly evident when compared to the oviposited group maintained under low nutrient conditions (Fig. 3). Increased mortality in oviposited mosquitoes maintained under low nutrient treatment, especially towards the later stages of the infection may account for the weak but nonetheless significant, negative effect of reduced dextrose on survival. In general, nutrition and oviposition status showed a more pronounced influence on the parasite traits of EIP and densities than on vector survival (Fig. 1A, Fig. 2A and Fig. 3A), which could be due to the reduced costs to vector survival at the lower temperature (20°C) [61] necessitated by this parasite-vector interaction [49]. Taken together, while the results presented here are in line with evolutionary theory for insects in general, they also indicate that the benefits of nutrient resorption to future fitness may not only outweigh the costs of reduced fecundity to the mosquitoes, while also enhancing their capacity to vector disease.

Follicular resorption has been suggested to result in reduced fecundity of *An. stephensi* following infections with another rodent malaria species, *P. yoelii nigeriensis* [60], and *An. maculipennis* infected with microsporidia [62]. To the best of the authors’ knowledge, whether follicular resorption directly benefits parasite replication has not been assessed empirically but is nevertheless plausible, and likely warrants confirmation. For instance, while greater availability of neutral lipids during oogenesis enhanced rates of sporozoite replication and migration to the salivary glands ( [63] and reviewed by [64]), similar lipid profiles were recovered following follicular resorption in *Ae. aegypti* [58]. Furthermore, in previous studies with the major arbovirus vector *Ae. aegypti*, higher rates of follicular resorption were observed in mosquitoes provided lower concentrations of sugar, likely to compensate for the reduced nutritional availability (at the cost of reduced fecundity) [57, 58]. In principle, higher rates of resorption in non-oviposited *An. stephensi* maintained in a low nutrition environment (1% dextrose) could explain the earlier appearance of *P. berghei* sporozoites in the salivary glands (yellow line, right pane, Fig. 1A), while continued availability of higher nutrient concentrations (10% dextrose + follicular resorption) in the other non-oviposited group could explain the higher proportion of infected mosquitoes over time, as a result of sustained sporozoite replication and/or migration (blue line, right pane, Fig. 1A) (and possibly enhanced survival, right pane, Fig. 3A) [49, 50]. Yet, when considered in the context of their oviposited counterparts, where a linear effect of dextrose on parasite fitness was evident in oviposited mosquitoes (shortened EIP and higher rates of infection, vector survival and sporozoite densities), it is unclear why the same 10-fold higher concentration of dextrose (and additional nutrients, albeit low, available from follicle resorption) in non-oviposited mosquitoes did not enhance rates of sporozoite replication and/or migration to the salivary gland, or support higher sporozoite densities. Taken together, these results suggest parasite fitness may be dependent on complex, dynamic interactions between physiological (nutrition) and evolutionary (egg retention) trade-offs in the mosquito vector.

## CONCLUSIONS

In summary, as with other insects, preventing mosquitoes from ovipositing (and completing their gonotrophic cycle) enhanced survival, even under nutrient limiting conditions [25, 32], albeit at the cost of promoting fitness of parasite species, whose analogs in humans cause malaria [43]. While increasing nutrition (% dextrose) positively impacted parasite fitness in mosquitoes that had laid eggs, the results from non-oviposited mosquitoes were consistent with the rates of follicular resorption compensating for differences in nutritional availability, and in the process enhancing vectorial capacity due to higher vector survival and earlier sporozoite appearance in the salivary glands (shorter EIP) as well as sporozoite densities. By obtaining three critical measures of transmission potential [45, 46, 47], this study argues for careful consideration of egg retention and nutritional availability on measures of vectorial capacity and vector competence.

From the parasite’s perspective, the overall patterns indicate an intrinsic ability to adjust its growth rates to the nutritional status of the host. Future studies manipulating the length of the gonotrophic cycle and duration of nutritional availability should reveal the extent of parasite’s adaptive plasticity. Additionally, from the vector’s perspective, whether parasite infection and low nutrition reduce fecundity and/or hatchability would be valuable to determine its effect on vector abundance, although neither trait appears to be as critical as selective oviposition behavior, which prioritizes egg retention [25, 27, 30, 31, 32, 34, 35, 58]. In general, however, as these results suggest, the consequences of the interactions between physiological (nutrition) and evolutionary (egg retention) trade-offs in the vector are likely to be complex for parasite fitness. As such, this complexity further underscores the importance of quantifying the contribution of selective oviposition behavior to disease transmission [41, 65], especially since most integrated vector control strategies [1, 14, 15, 16, 17, 18, 19] and human-induced climate change either directly or indirectly target this key mosquito behavior [20, 21, 22, 23, 24].

## LEGENDS TO SUPPLEMENTARY FIGURES AND TABLES

**Additional file 1: Fig. S1. A)** Study design. **B**) Representative images of ovaries from mosquitoes that had oviposited successfully (left pane) or still retained eggs (right pane).

**Additional file 2: Fig. S2. A)** The proportion of mosquitoes that oviposited after being provided an oviposition site (left pane, referred to as ‘oviposited’ elsewhere in this manuscript) or not provided a site (right pane, referred to as ‘non-oviposited’ elsewhere), based on the gravid status (presence or absence of eggs as in Additional file 1: Fig. S1B) of mosquitoes at the time of dissection. **B)** Mean rates of gravidity recorded at the time points in A). Note that the data in this figure is representative of all 616 mosquitoes sampled in the study (Fig. 1 and Fig. 2) as the same approach was used to assess gravid status, prior to preparing the salivary glands to assess prevalence (Fig. 1) or densities (Fig. 2). For statistical analysis of the data, refer to Additional file 3: Table S1.

**Additional file 3: Table S1.** Statistical models of the proportion of mosquitoes that had oviposited after being provided an oviposition site (‘oviposited’) or not provided a site (‘non-oviposited’), based on the gravid status (‘*y* = gravid status (ovaries ± eggs)’) at the time of dissections for data collection, as depicted in Additional file 1: Fig. S1.

**Additional file 4: Figure S3.** Excluding individuals that did not oviposit when offered an oviposition site, or oviposited despite not being offered an oviposition sites, does not influence the mean rates of sporozoite prevalence (**A**) or densities (**B**). Each data point represents the mean of mosquitoes sampled from all four groups at each time point (for prevalence, 4 groups * 7 dpi = 28 data points, for densities, 4 groups * 5 dpi = 20 data points); x-axes in both figures depicting the mean of all individuals (regardless of oviposition status), regressed against the mean of individual mosquitoes that were confirmed to have oviposited when provided a site (‘oviposited’), or gravid (carrying eggs) due to not being provided an oviposition site (‘non-oviposited’).

**Additional file 5: Table S2**. Pairwise comparisons (post-hoc) of the estimated marginal means in sporozoite prevalence predicted by the statistical model in Table 1 (‘*y* = sporozoite prevalence’) (Fig. 1A). Comparisons were made between each of the four groups (oviposited and nutrient treatments) for the indicated day post-infection. Estimates (‘Est. (= A-B)’) indicate the difference of the predicted mean in group B subtracted from that of group A. Tukey’s method was used to adjust for multiple comparisons.

**Additional file 6: Table S3**. Pairwise comparisons (post-hoc) of the estimated marginal means in sporozoite densities predicted by the statistical model in Table 1 (‘*y* = sporozoite densities’) (Fig. 2A). Comparisons were made between each of the four groups (oviposited and nutrient treatments) for the indicated day post-infection. Estimates (‘Est. (= A-B)’) indicate the difference of the predicted mean in group B, subtracted from that of group A. Tukey’s method was used to adjust for multiple comparisons.

**Additional file 7: Table S4**. Statistical model of survival rates (Fig. 3). Survival was measured daily in all four groups until 27 dpi, and data modeled using Cox proportional hazards.

## ETHICS APPROVAL AND CONSENT TO PARTICIPATE

Not applicable.

## CONSENT FOR PUBLICATION

Not applicable

## AVAILABILITY OF DATA AND MATERIALS

The datasets used and/or analysed during the current study are available from the corresponding author on reasonable request.

## AUTHORS’ CONTRIBUTIONS

JCS: Design of the work, acquisition, analysis, interpretation, drafted the work.

NGD: acquisition, analysis.

DEK: acquisition.

AKP: Conception, design of the work, acquisition, analysis, interpretation, drafted the work and substantially revised.

All authors have approved the submitted version.

## ACKNOWLEDGEMENTS

The authors would like to thank Courtnie Vickery and Rafael Freitas for help with data collection, Anne Elliot and Jared Cooper for their help with mosquito husbandry, and the staff at UGA animal resources for mouse husbandry.

