## Additional file 1: Figure S1 for "Oviposition status and sugar availability influence measures of transmission potential in malaria-infected mosquitoes"

**A.**

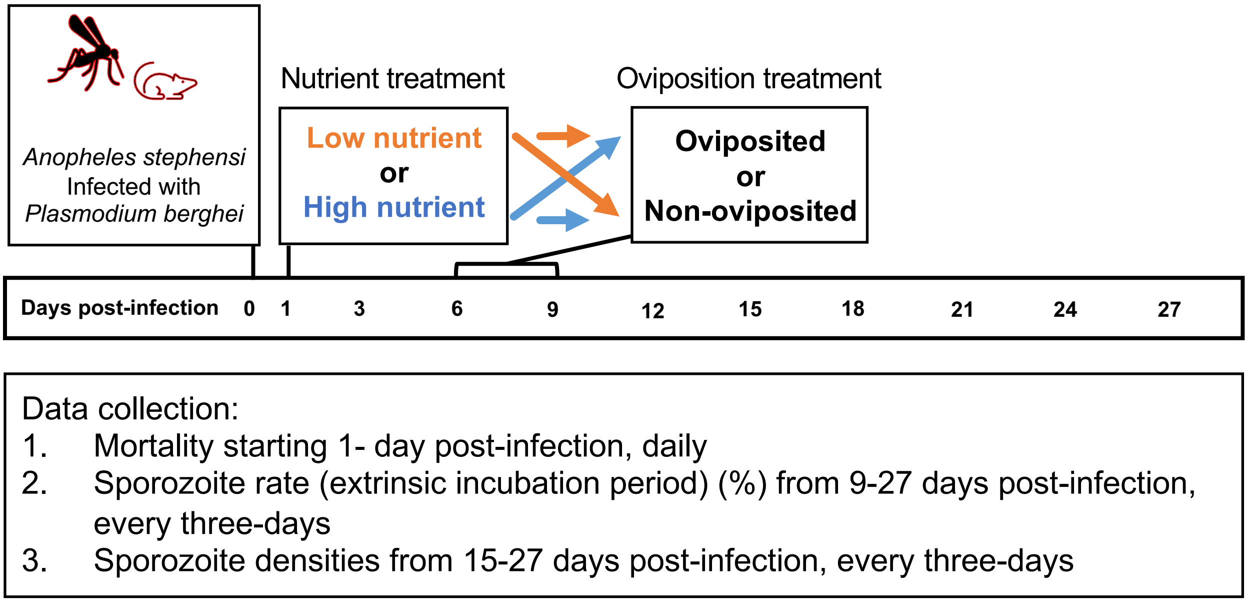


**B.**

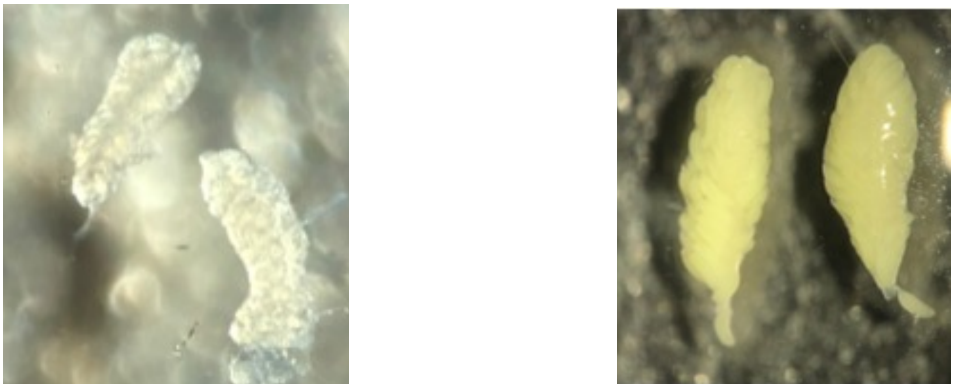

Ovaries from oviposited individuals

Ovaries from non-oviposited individuals
