## Supplementary figures and images for "Oviposition status and sugar availability influence measures of transmission potential in malaria-infected mosquitoes"

### Additional file 2: Figure S2

**Additional file 2: Figure S2**

A.

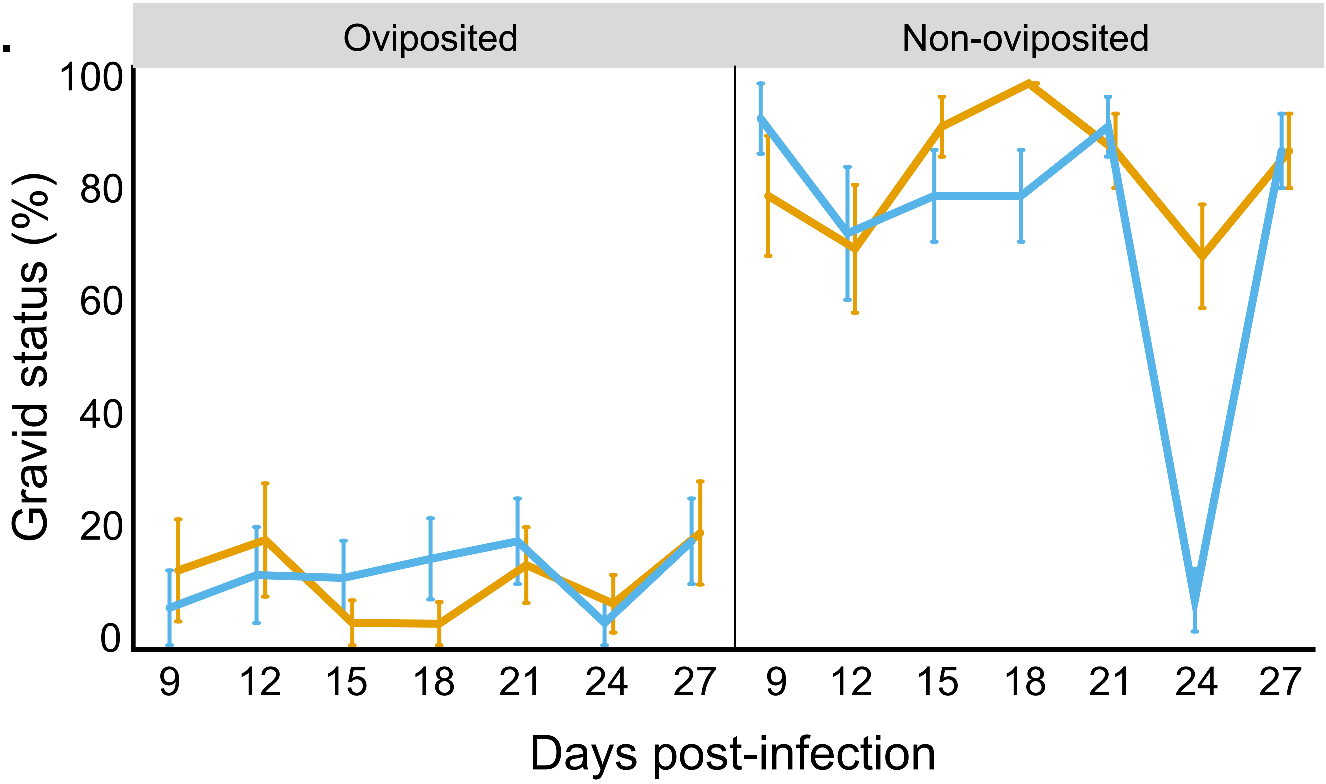


B.

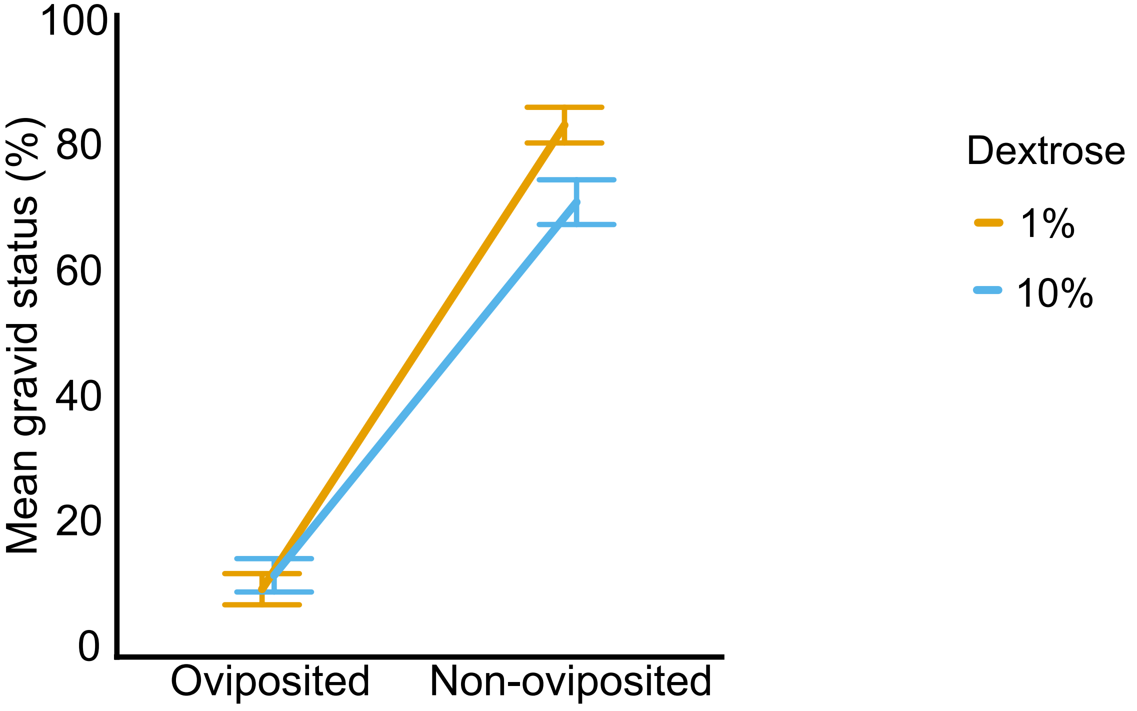

### Additional file 4: Figure S3

#### Additional file 4: Figure S3

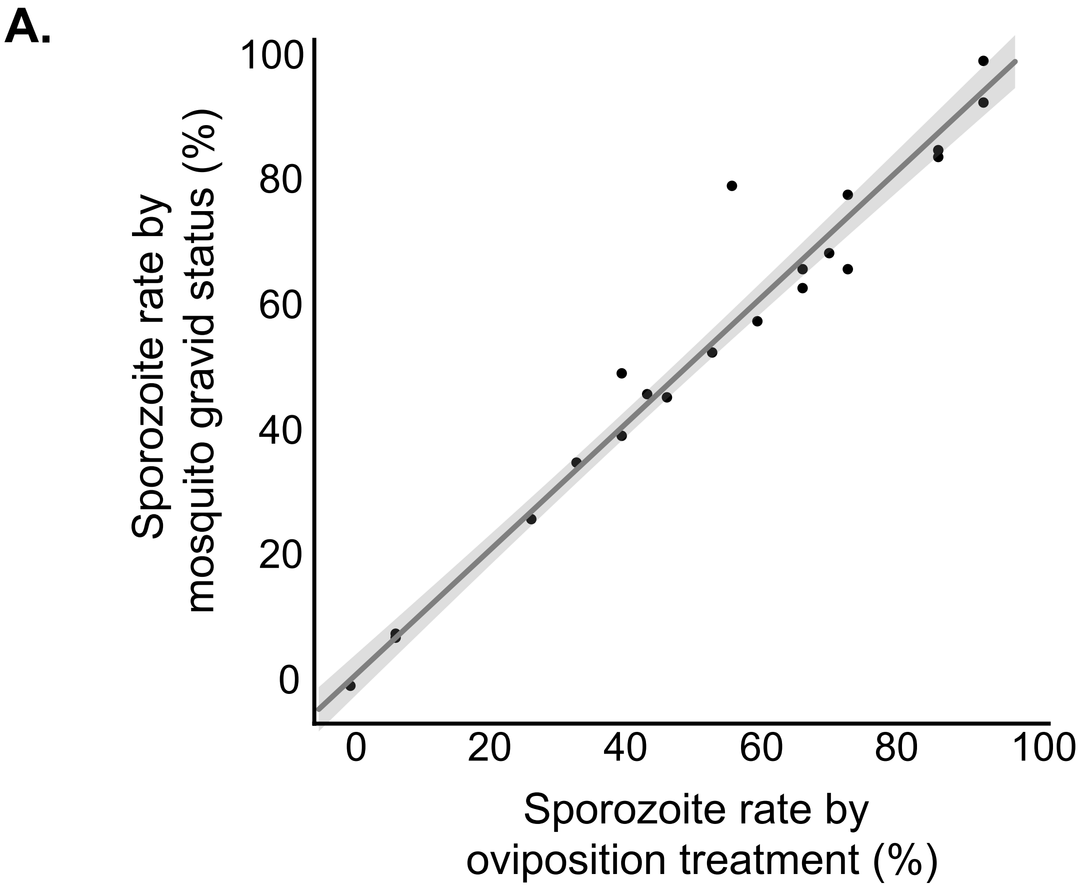


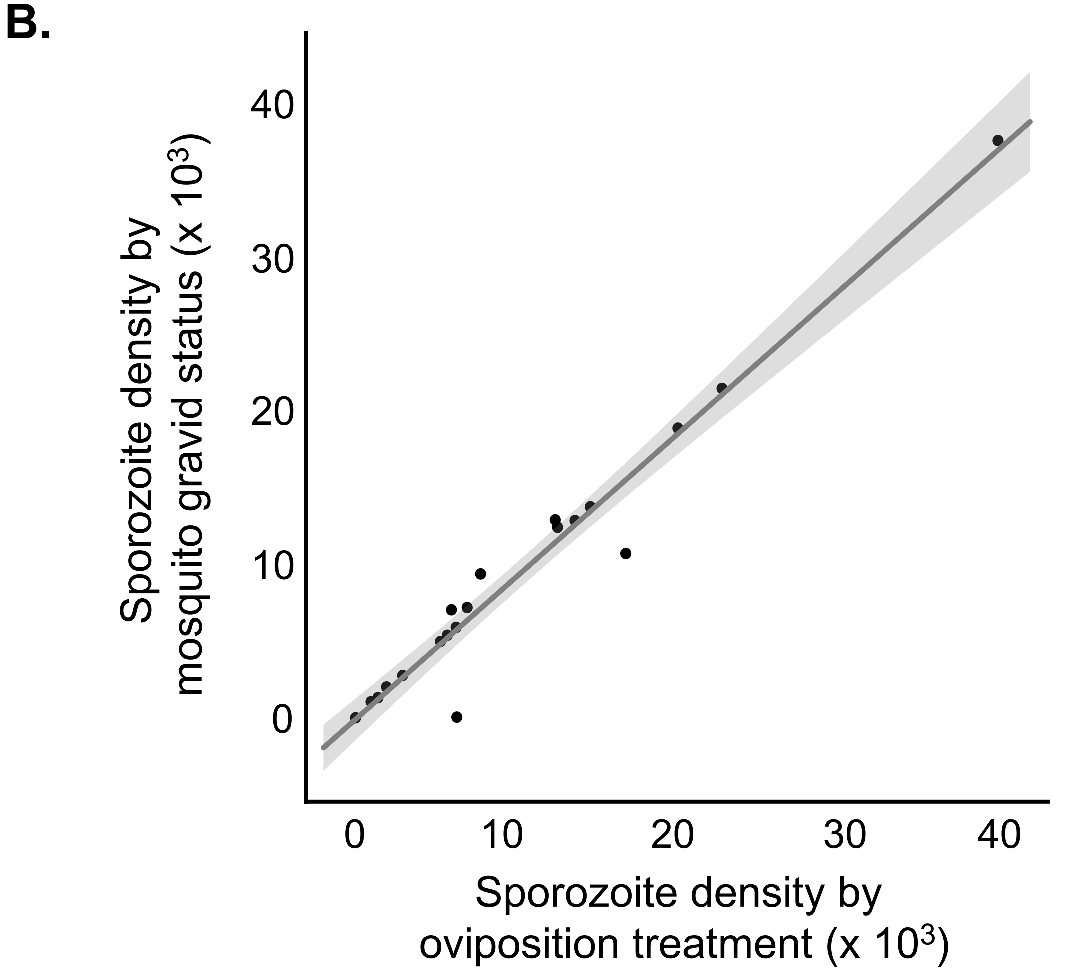
