## Additional file 3: Table S1 for "Oviposition status and sugar availability influence measures of transmission potential in malaria-infected mosquitoes"

| **Table S1** | | | | | | | |
| --- | --- | --- | --- | --- | --- | --- | --- |
| *y* = gravid status (ovaries ± eggs) | | Oviposited ^a1^ | | | Non-oviposited ^a2^ | | |
| Row | Terms (*x*) | *χ^2^* | *df* | *P* | *χ^2^* | *df* | *P* |
| 2 | Nutrient levels ^b^ | 0.49 | 1 | 0.484 | 6.86 | 1 | 0.009* |
| 3 | Dpi ^c^ (linear effect) | 0.02 | 1 | 0.884 | 3.38 | 1 | 0.066 |
| 7 | Nutrient levels * dpi | 0.22 | 1 | 0.638 | 2.75 | 1 | 0.097 |
|  | Number of mosquito ovaries assessed | 304 | | | 312 | | |
| ^a1^ Oviposited = mosquitoes provided oviposition sites; ^a2^ non-oviposited = mosquitoes not provided oviposition sites; ^b^ 1 or 10% dextrose; ^c^ salivary glands sampled at 3-day intervals from 9-27 dpi. | | | | | | | |
| Abbreviations:  *χ^2^*= Chi-squared test (type 2); *df* = degrees of freedom; dpi = days post-infection; *P* = p-value. | | | | | | | |
| * Statistically significant (i.e., *P*<0.05). | | | | | | | |
