## Additional file 5: Table S2 for "Oviposition status and sugar availability influence measures of transmission potential in malaria-infected mosquitoes"

| **Table S2** | | | | | | |
| --- | --- | --- | --- | --- | --- | --- |
| Row | A | B | dpi | Est. (=A-B) | SE | P |
| 1 | Oviposited, 1% dextrose | Non-oviposited, 1% dextrose | 9 | 0.00 | 0.00 | 0.783 |
| 2 | Oviposited, 1% dextrose | Oviposited, 10% dextrose | 9 | -0.01 | 0.01 | 0.704 |
| 3 | Oviposited, 1% dextrose | Non-oviposited, 10% dextrose | 9 | 0.00 | 0.00 | 0.975 |
| 4 | Oviposited, 1% dextrose | Non-oviposited, 1% dextrose | 12 | -0.04 | 0.03 | 0.433 |
| 5 | Oviposited, 1% dextrose | Oviposited, 10% dextrose | 12 | -0.09 | 0.05 | 0.226 |
| 6 | Oviposited, 1% dextrose | Non-oviposited, 10% dextrose | 12 | 0.00 | 0.01 | 0.999 |
| 7 | Oviposited, 1% dextrose | Non-oviposited, 1% dextrose | 15 | -0.23 | 0.09 | 0.046* |
| 8 | Oviposited, 1% dextrose | Oviposited, 10% dextrose | 15 | -0.37 | 0.10 | <0.001* |
| 9 | Oviposited, 1% dextrose | Non-oviposited, 10% dextrose | 15 | -0.03 | 0.06 | 0.958 |
| 10 | Oviposited, 1% dextrose | Non-oviposited, 1% dextrose | 18 | -0.36 | 0.10 | 0.002* |
| 11 | Oviposited, 1% dextrose | Oviposited, 10% dextrose | 18 | -0.49 | 0.09 | <0.001* |
| 12 | Oviposited, 1% dextrose | Non-oviposited, 10% dextrose | 18 | -0.21 | 0.12 | 0.245 |
| 13 | Oviposited, 1% dextrose | Non-oviposited, 1% dextrose | 21 | -0.32 | 0.08 | <0.001* |
| 14 | Oviposited, 1% dextrose | Oviposited, 10% dextrose | 21 | -0.40 | 0.08 | <0.001* |
| 15 | Oviposited, 1% dextrose | Non-oviposited, 10% dextrose | 21 | -0.34 | 0.09 | <0.001* |
| 16 | Oviposited, 1% dextrose | Non-oviposited, 1% dextrose | 24 | -0.29 | 0.11 | 0.043* |
| 17 | Oviposited, 1% dextrose | Oviposited, 10% dextrose | 24 | -0.37 | 0.10 | 0.002* |
| 18 | Oviposited, 1% dextrose | Non-oviposited, 10% dextrose | 24 | -0.42 | 0.10 | <0.001* |
| 19 | Oviposited, 1% dextrose | Non-oviposited, 1% dextrose | 27 | -0.25 | 0.15 | 0.364 |
| 20 | Oviposited, 1% dextrose | Oviposited, 10% dextrose | 27 | -0.35 | 0.15 | 0.096 |
| 21 | Oviposited, 1% dextrose | Non-oviposited, 10% dextrose | 27 | -0.55 | 0.14 | <0.001* |
| 22 | Non-oviposited, 1% dextrose | Oviposited, 10% dextrose | 9 | 0.00 | 0.00 | 0.901 |
| 23 | Non-oviposited, 1% dextrose | Non-oviposited, 10% dextrose | 9 | 0.00 | 0.00 | 0.762 |
| 24 | Non-oviposited, 1% dextrose | Oviposited, 10% dextrose | 12 | -0.04 | 0.05 | 0.794 |
| 25 | Non-oviposited, 1% dextrose | Non-oviposited, 10% dextrose | 12 | 0.04 | 0.03 | 0.404 |
| 26 | Non-oviposited, 1% dextrose | Oviposited, 10% dextrose | 15 | -0.14 | 0.12 | 0.625 |
| 27 | Non-oviposited, 1% dextrose | Non-oviposited, 10% dextrose | 15 | 0.20 | 0.09 | 0.141 |
| 28 | Non-oviposited, 1% dextrose | Oviposited, 10% dextrose | 18 | -0.12 | 0.08 | 0.466 |
| 29 | Non-oviposited, 1% dextrose | Non-oviposited, 10% dextrose | 18 | 0.15 | 0.11 | 0.536 |
| 30 | Non-oviposited, 1% dextrose | Oviposited, 10% dextrose | 21 | -0.08 | 0.06 | 0.468 |
| 31 | Non-oviposited, 1% dextrose | Non-oviposited, 10% dextrose | 21 | -0.02 | 0.07 | 0.992 |
| 32 | Non-oviposited, 1% dextrose | Oviposited, 10% dextrose | 24 | -0.08 | 0.07 | 0.709 |
| 33 | Non-oviposited, 1% dextrose | Non-oviposited, 10% dextrose | 24 | -0.13 | 0.07 | 0.3 |
| 34 | Non-oviposited, 1% dextrose | Oviposited, 10% dextrose | 27 | -0.10 | 0.14 | 0.895 |
| 35 | Non-oviposited, 1% dextrose | Non-oviposited, 10% dextrose | 27 | -0.30 | 0.12 | 0.072 |
| 36 | Oviposited, 10% dextrose | Non-oviposited, 10% dextrose | 9 | 0.01 | 0.01 | 0.695 |
| 37 | Oviposited, 10% dextrose | Non-oviposited, 10% dextrose | 12 | 0.09 | 0.05 | 0.215 |
| 38 | Oviposited, 10% dextrose | Non-oviposited, 10% dextrose | 15 | 0.34 | 0.10 | 0.004* |
| 39 | Oviposited, 10% dextrose | Non-oviposited, 10% dextrose | 18 | 0.27 | 0.10 | 0.048* |
| 40 | Oviposited, 10% dextrose | Non-oviposited, 10% dextrose | 21 | 0.06 | 0.06 | 0.747 |
| 41 | Oviposited, 10% dextrose | Non-oviposited, 10% dextrose | 24 | -0.05 | 0.07 | 0.875 |
| 42 | Oviposited, 10% dextrose | Non-oviposited, 10% dextrose | 27 | -0.21 | 0.12 | 0.315 |
