## Additional file 6: Table S3 for "Oviposition status and sugar availability influence measures of transmission potential in malaria-infected mosquitoes"

| **Table S3** | | | | | | |
| --- | --- | --- | --- | --- | --- | --- |
| Row | A | B | dpi | Est. (=A-B) | SE | P |
| 1 | Oviposited, 1% dextrose | Non-oviposited, 1% dextrose | 15 | 968.73 | 1440.58 | 0.908 |
| 2 | Oviposited, 1% dextrose | Oviposited, 10% dextrose | 15 | -5871.40 | 3819.16 | 0.415 |
| 3 | Oviposited, 1% dextrose | Non-oviposited, 10% dextrose | 15 | -2772.36 | 2521.66 | 0.69 |
| 4 | Oviposited, 1% dextrose | Non-oviposited, 1% dextrose | 18 | -378.30 | 1718.59 | 0.996 |
| 5 | Oviposited, 1% dextrose | Oviposited, 10% dextrose | 18 | -10212.81 | 3955.95 | 0.048* |
| 6 | Oviposited, 1% dextrose | Non-oviposited, 10% dextrose | 18 | -3538.90 | 2327.30 | 0.425 |
| 7 | Oviposited, 1% dextrose | Non-oviposited, 1% dextrose | 21 | -3920.06 | 2132.94 | 0.256 |
| 8 | Oviposited, 1% dextrose | Oviposited, 10% dextrose | 21 | -14819.56 | 4172.84 | 0.002* |
| 9 | Oviposited, 1% dextrose | Non-oviposited, 10% dextrose | 21 | -3790.14 | 2068.61 | 0.258 |
| 10 | Oviposited, 1% dextrose | Non-oviposited, 1% dextrose | 24 | -9757.14 | 3719.25 | 0.043* |
| 11 | Oviposited, 1% dextrose | Oviposited, 10% dextrose | 24 | -17985.16 | 5761.23 | 0.01* |
| 12 | Oviposited, 1% dextrose | Non-oviposited, 10% dextrose | 24 | -3402.19 | 2269.61 | 0.438 |
| 13 | Oviposited, 1% dextrose | Non-oviposited, 1% dextrose | 27 | -16504.40 | 7624.68 | 0.133 |
| 14 | Oviposited, 1% dextrose | Oviposited, 10% dextrose | 27 | -18288.80 | 8840.04 | 0.163 |
| 15 | Oviposited, 1% dextrose | Non-oviposited, 10% dextrose | 27 | -2556.16 | 2624.10 | 0.764 |
| 16 | Non-oviposited, 1% dextrose | Oviposited, 10% dextrose | 15 | -6840.13 | 3688.47 | 0.248 |
| 17 | Non-oviposited, 1% dextrose | Non-oviposited, 10% dextrose | 15 | -3741.09 | 2302.08 | 0.364 |
| 18 | Non-oviposited, 1% dextrose | Oviposited, 10% dextrose | 18 | -9834.51 | 3881.58 | 0.055 |
| 19 | Non-oviposited, 1% dextrose | Non-oviposited, 10% dextrose | 18 | -3160.60 | 2193.80 | 0.474 |
| 20 | Non-oviposited, 1% dextrose | Oviposited, 10% dextrose | 21 | -10899.50 | 4288.35 | 0.054 |
| 21 | Non-oviposited, 1% dextrose | Non-oviposited, 10% dextrose | 21 | 129.92 | 2403.12 | 1 |
| 22 | Non-oviposited, 1% dextrose | Oviposited, 10% dextrose | 24 | -8228.03 | 6612.97 | 0.599 |
| 23 | Non-oviposited, 1% dextrose | Non-oviposited, 10% dextrose | 24 | 6354.95 | 3976.84 | 0.38 |
| 24 | Non-oviposited, 1% dextrose | Oviposited, 10% dextrose | 27 | -1784.41 | 10800.06 | 0.998 |
| 25 | Non-oviposited, 1% dextrose | Non-oviposited, 10% dextrose | 27 | 13948.24 | 7717.57 | 0.27 |
| 26 | Oviposited, 10% dextrose | Non-oviposited, 10% dextrose | 15 | 3099.04 | 3963.81 | 0.863 |
| 27 | Oviposited, 10% dextrose | Non-oviposited, 10% dextrose | 18 | 6673.91 | 4185.10 | 0.382 |
| 28 | Oviposited, 10% dextrose | Non-oviposited, 10% dextrose | 21 | 11029.42 | 4285.26 | 0.049* |
| 29 | Oviposited, 10% dextrose | Non-oviposited, 10% dextrose | 24 | 14582.98 | 5929.55 | 0.066 |
| 30 | Oviposited, 10% dextrose | Non-oviposited, 10% dextrose | 27 | 15732.64 | 8910.91 | 0.29 |
