## Additional file 7: Table S4 for "Oviposition status and sugar availability influence measures of transmission potential in malaria-infected mosquitoes"

| **Table S4** | | | | |
| --- | --- | --- | --- | --- |
|  |  | *y* = daily mortality risk | | |
| Row | Terms (*x*) | *χ^2^* | *df* | *P* |
| 1 | Oviposition status ^a^ | 15.37 | 1 | <0.001* |
| 2 | Nutrient levels ^b^ | 4.94 | 1 | 0.026* |
| 3 | Oviposition status * nutrient levels | 2.36 | 1 | 0.124 |
| ^a^ Oviposited or Non-oviposited mosquitoes; ^b^ 1 or 10% dextrose. | | | | |
| Abbreviations:  *χ^2^*= Chi-squared test (type 2); *df* = degrees of freedom; *P* = p-value. | | | | |
